# Myeloperoxidase promotes fibrosis by inhibiting cathepsin K to bias the lung toward ECM accumulation

**DOI:** 10.64898/2026.04.05.713467

**Authors:** Patrick A. Link, Jack Wellmerling, Jeffrey A. Meridew, Hyogo Naoi, YS Prakash, Mauricio Rojas, Eva M. Carmona, Daniel J. Tschumperlin

## Abstract

Pulmonary fibrosis (PF) involves excessive collagen accumulation, yet mechanisms shifting the balance of synthesis and degradation toward net deposition remain unclear. Myeloperoxidase (MPO) inversely correlates with survival in PF. Using the bleomycin model, we found MPO knockout (MPOko) mice were protected from fibrosis, and pharmacological MPO inhibition after peak inflammation (day 7) recapitulated this protection. MPO persisted in lung tissue 21 days post-injury despite neutrophil efflux, linking acute inflammation to sustained remodeling. Mechanistically, we identified that MPO inhibits Cathepsin K (CatK), a potent collagenolytic enzyme involved in fibrosis resolution. Notably, CatK gene expression (*CTSK*) is elevated in PF, suggesting post-translational inhibition of CatK. MPOko and inhibitor-treated mice exhibited elevated CatK activity after bleomycin; exogenous addition of pathophysiologic concentrations of MPO reduced CatK activity in mouse precision-cut lung slices and human fibroblasts. Biochemically, MPO reduced CatK activity to 33% of control. In two distinct cohorts of PF patients, we observed significantly increased MPO protein levels in platelet poor plasma and in lung tissue. In PF patients, 62% had MPO levels in platelet poor plasma exceeding healthy controls, while lung tissue from other PF patients showed significantly elevated MPO staining. Plasma levels were inversely correlated with decreased survival, FVC, and DLCO. These findings establish MPO as a post-translational inhibitor of CatK-mediated collagenolysis, revealing a mechanism linking acute inflammation to sustained fibrosis and suggest a patient subpopulation that may benefit from MPO-targeted therapy.

**Graphical Abstract:** 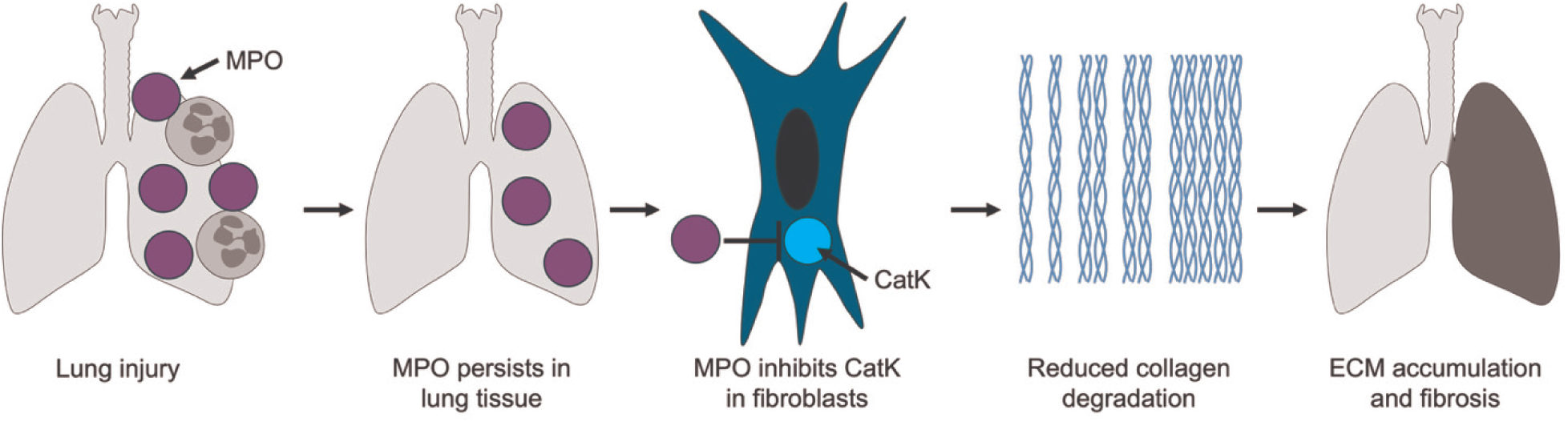

Myeloperoxidase persists in lung tissue after injury and inhibits cathepsin K activity, impairing collagen degradation and promoting extracellular matrix accumulation during pulmonary fibrosis.

## Introduction

In pulmonary fibrosis (PF), tissue remodeling and interstitial scarring stiffens the lungs, leading to respiratory failure. Scarring develops through excessive accumulation of extracellular matrix (ECM), the structural scaffolding that supports lung cell and tissue functions. The accumulation of ECM can occur through multiple mechanisms including increased deposition or decreased resorption. During homeostasis, collagen in the lung is continuously deposited and resorbed,^1–5^ but in PF, a shift toward increased deposition and decreased resorption has been observed.^6,7^ Despite advances in understanding ECM deposition, the factors that prevent normal ECM resorption remain poorly characterized.^3,6–15^

Fibroblasts, the primary cells responsible for matrix maintenance in healthy tissue, are a key therapeutic target to restore the ECM accumulation balance. To degrade collagen, fibroblasts produce collagenolytic enzymes like matrix metalloproteases and cathepsins. Cathepsin K (CatK), a cysteine protease, is unique among cathepsins in its ability to cleave the collagen triple helix at multiple sites.^16,17^ We and others have shown the importance of CatK in pulmonary fibrosis, with overexpression of CatK reducing fibrosis and increasing collagen degradation, and CatK knockout increasing fibrosis.^18–20^ Although increased CatK gene expression has been suggested as a negative prognostic indicator in patients with IPF,^21^ expression does not necessarily reflect activity. CatK must be activated from pro-CatK before it can break down collagens, and once activated, CatK activity can be attenuated by reactive oxygen species (ROS).^22^ ROS are dramatically increased in IPF,^23,24^ suggesting that the oxidative environment may impair collagenolytic activity, even in the presence of increased CatK expression.

Injury and inflammation precede fibrosis in human disease and animal models and infiltrating inflammatory cells represent a major source of extracellular ROS in the injured lung. Neutrophils are among the first responders to injury, and elevated neutrophil counts predict worse outcomes in PF patients.^5,25–31^ In acute exacerbations of IPF (AE-IPF), a well characterized neutrophil surge exists where the fibrotic progression occurs in weeks rather than years, and accounts for ∼46% of IPF mortality ^32–35^. However, in stable IPF patients, neutrophilia is minimal,^36^ suggesting initial infiltration after injury and a return to baseline. This is supported by animal models of fibrosis that show increased neutrophil infiltration with the initial injury that returns to baseline while fibrosis continues. This temporal disconnect has led the field to largely exclude neutrophils as significant contributors to the ECM accumulation that occurs in PF. This view is further supported by evidence that anti-inflammatory therapies can be harmful in PF, while broad neutrophil depletion in animal models produces ambiguous results.^37–40^ However, in other contexts, neutrophils participate in complex ECM regulatory networks,^41–45^ raising the possibility that specific neutrophil-derived factors, rather than neutrophils themselves, may influence fibrosis progression.

Myeloperoxidase (MPO) represents a candidate link between inflammatory injury and sustained ECM accumulation. When activated, neutrophils release MPO into surrounding tissue, where it catalyzes conversion of hydrogen peroxide into highly reactive oxidative species as a primary component of innate immunity. Because of its role in generating these highly reactive species, MPO has long been associated with acute lung injury and adult respiratory distress syndrome. However, tissue-bound MPO persists long after neutrophil efflux and retains activity,^46–48^ possibly acting as a temporal bridge connecting transient inflammation to sustained fibrosis. Importantly MPO has emerged as a mediator of fibrosis in multiple organ systems; MPO-deficient (MPOko)mice are protected from atrial, renal, and liver fibrosis, while pharmacological MPO inhibition reduces cardiac remodeling after myocardial infarction.^49–51^ In patients with IPF, high MPO levels inversely correlate with survival and vital capacity.^23,31,52^ Unbiased proteomic analyses have identified elevated MPO in the fibrotic alveoli of IPF patients and in bronchoalveolar lavage fluid of other interstitial lung diseases.^31,53,54^ Despite MPO’s established profibrotic role in other organs and its clinical association with IPF severity, whether MPO directly contributes to pulmonary fibrosis and collagen turnover by lung cells has not been tested.

Based on these observations, we hypothesized that MPO inhibits CatK-mediated collagen degradation, providing a mechanism by which neutrophil-derived factors promote fibrosis. In this study, we demonstrate that MPOko and pharmacological MPO inhibition protect against bleomycin-induced fibrosis in mice, with protection evident even when inhibition begins after the peak of the inflammatory phase. We identify CatK inhibition as a mechanism underlying this effect, showing that MPO directly suppresses CatK activity in recombinant systems, fibroblasts, and lung tissue. We further show that MPO persists in lung tissue after neutrophils have returned to normal levels, and that a subset of IPF patients exhibit elevated circulating and tissue MPO levels. These findings establish MPO as a regulator of collagen degradation and suggest that inhibiting MPO could partially restore ECM homeostasis in pulmonary fibrosis.

## Methods

### Study Approvals

All non-human animal procedures were performed in accordance with Mayo’s Institutional Animal Care and Use Committee. All efforts were made to minimize animal suffering and reduce the number of animals used. Mice used in this study were housed in standard laboratory conditions with a 12:12 hour light-dark cycle, temperature controlled, and relative humidity controlled. Food and water were provided ad libitum and animals were monitored daily for health and well-being.

All procedures performed in this study involving human participants were in accordance with the ethical standards of the Institutional Review Board (OSU IRB#2019H0310, Mayo Clinic IRBs #14-005305 and #17-008088). Informed consent was obtained from all individual participants included in the study. Patient demographics can be found in the Supplement (Supp. Table 1).

### Sex as a Biological Variable

Our study examined male and female human subjects. Patient demographics can be found in the Supplement (Supp. Table 1). Approximately 67% of human samples were from males, and sex-dimorphic effects are shown.

Our study examined male and female mice, and similar findings are reported for both sexes.

### Reagents

Comprehensive reagent details can be found in Supplement (Supp. Table 2).

### Bleomycin-Induced Fibrosis Model

To establish pulmonary fibrosis in 8–12-week-old WT (C57Bl/6J) or MPOko mice, we anesthetized mice with isoflurane. Following anesthesia, we intubated each mouse using a 20-gauge catheter (BD angiocath #26742) and confirmed proper tracheal placement through PBS bubble visualization as previously described.^55^ 50 µl PBS (vehicle control) or bleomycin (1 U/kg, Hospira Inc. in PBS) was delivered intratracheally. Survival was tracked daily and weights were taken weekly.

At indicated timepoints, mice were anesthetized with Ketamine and Xylazine (100/10 mg/kg), euthanized, and blood was collected through cardiac puncture. Intratracheal catheterization was performed, and 1 mL (500 µL two times) cold PBS was administered and removed to collect BALF. We then perfused the right ventricle with ice-cold PBS containing 2 mM EDTA. Lungs used for histology or downstream tissue analysis.

To quantify collagen content, hydroxyproline was measured using a hydroxyproline assay kit (BioVision K555) based on manufacturer’s instructions.

To quantify histology, sections (5 μm thick) were cut from formalin-fixed paraffin-embedded lung tissues, and the sections were stained for Hematoxylin and Eosin (H&E), Masson’s Trichrome (MTC), or picrosirius red (PSR). Brightfield images were taken with a Motic EasyScan for H&E or MTC while fluorescent images were taken with a Zeiss Axio Observer. Whole lung images were analyzed as previously described for injury level (H&E),^56^ fibrotic regions (MTC),^57^ and collagen detection (PSR).^58^

For flow cytometry, tissue dissociation was performed as previously described with minor changes.^55^ We harvested lungs and minced them in 10 cm Petri dishes separating a portion for flow cytometry before enzymatic digestion. The tissue underwent digestion in DMEM containing Liberase TM (0.1 mg/mL) and DNase I (100 U/mL) for 35 minutes at 37°C with constant rotation. We halted digestion with DMEM containing 10% FBS, then filtered the suspension through 40 µm mesh to remove debris. After centrifugation and RBC lysis, we added surface antibodies to stain various populations of cells including: CD3, CD11c, CD11b, CD64, Ly-6G, CD170, CD45, CD31, Ly-6C, and viability stain. We performed flow cytometry on a BD LSR Fortessa X-20 (gating strategy shown in Supp. Fig. 1) and analyzed the results using FlowJo v10.

Flow cytometry of BALF and blood was performed as described above without tissue dissociation. Blood was additionally processed with Lysis buffer at room temperature for 15 minutes before antibody staining.

To quantify MPO in tissue, lung tissue snap frozen in liquid nitrogen and store at −80°C. Frozen tissue was homogenized using a BeadMill 24 and 2 mm ceramic beads on the Lung tissue setting in T-PER. Homogenized tissue was centrifuged at 10,000 g for 5 minutes and the supernatant was used for the ELISA (R&D systems, DY3557) according to manufacturer’s directions.

### Reanalysis of Publicly Available single cell RNA sequencing data sets

Single cell RNA-sequencing datasets were analyzed using the depositing authors’ processing and annotations where available (GSE210341 and LMEX0000004396).^59,60^ For GSE132771, we filtered cells according to library size, number of expressed genes, and mitochondrial gene fraction according to the authors.^61^ Data from each donor were integrated using Seurat v3’s integration workflow.^62^ Cell types were annotated using Seurat’s reference-based annotation with mesenchymal cells from the Human Lung Cell Atlas. Figures were made using R.

### In vivo CatK Activity

CatK activity was quantified in vivo using IVISense Cat K 680 Fast Fluorescent Probe (Perkin Elmer, NEV11000). The 100 µL of the reconstituted probe was injected intravenously 13 days after bleomycin administration. 24 hours later, mice were anesthetized using isoflurane, shaved, and imaged using an In Vivo Imaging System (Perkin Elmer).

### Precision Cut Lung Slices (PCLS)

C57BL/6J mice aged from 8-12 weeks old were euthanized and the lungs were perfused with PBS containing 2 mM EDTA. We filled the lung airspaces with 10% gelatin as per ^63–65^ and submerged them in cold HBSS containing 10 mM HEPES 3 mM CaCl2, and 1 mM MgSO4. One mm of the base of the left lobe was cut off to increase adhesive surface area. The lungs were then glued to the piston of a Precisionary Vibratome (VF-210-0Z), embedded in 3% low melting point agarose, and sliced to 300 µm thickness.

PCLS were cultured for 4 days (media refreshed after 2 days) total in a 1:1 mixture of Eagle’s Minimum Essential Media (EMEM, Lonza) to Alveolar Epithelial Cell Media (ScienCell) supplemented with 6% FBS and 1% anti-anti. For collagen degradation assays, PCLS were cultured with MPO (0.2 ug/ml final concentration) or vehicle. Conditioned media was collected, and used to measure collagen synthesis and degradation products, Procollagen I N-terminal Propeptide (PINP) and C-telopeptide of type I collagen (CTX-I), respectively, using ELISA assays according to manufacturer’s directions and diluting the supernatant 1:2 in sample buffer.

To measure CatK activity, PCLS were cultured as described above with TGF-β (2 ng/ml final concentration), MPO (0.2 ug/ml final concentration), and/or 4-Aminobenzoic Acid Hydrazide (ABAH, 2.2 µM final concentration) or vehicle added to media for 4 days with media refreshed after 2 days. On the final day Magic Red (ImmunoChemistry, 940) was added to wells for 1 hour and then CatK activity was analyzed using a fluorescent plate reader (ex. 592, em. 628, cutoff 630) according to manufacturer’s directions.

### Cell Culture and Treatments

IMR-90 cells were cultured to confluence, then media was changed to contain 0.1% FBS, 20 ug/mL ascorbic acid and 2 ng/ml TGF-β for 6 days total, with media refreshed on day 3. On the fourth day H_2_O_2_ (450 µM; all wells), MPO (0.2 ug/ml) or vehicle was added to wells. On the sixth day Magic Red (ImmunoChemistry, 940) was added to wells for 1 hour and then analyzed using a fluorescent plate reader (ex 592, em 628, cutoff 630) according to manufacturer’s directions.

### Direct Biochemical Protein Experiments

Direct interaction assays were performed as previously described with modifications.^22^ In brief, Human CatK (2 nM) in the presence of 0.05 M sodium acetate buffer, pH 6.5, 100 µM NaCl (final concentration), was combined with MPO (0.2 ug/ml, from three different MPO sources). Then increasing concentrations of H_2_O_2_ were added to inactivate the CatK as previously described.^22^ Finally, Magic Red CatK activity probe was added and incubated at 37°C for 1 hour before being read on a fluorescent plate reader as per manufacturers recommendations. Substrate only (Magic Red), substrate plus treatment compounds (MPO or H2O2), activator only (CatK), and buffer only controls to verify changes in fluorescent activity reflect enzyme-dependent activation of Magic Red can be found in (Supp. Fig. 2).

### Human Plasma levels of MPO

Blood samples were collected under IRB#17-008088 approved protocol. Participants were recruited from the outpatient Interstitial Lung Disease (ILD) clinic and enrolled in the institutional biobank after providing written informed consent. Eligible participants had a diagnosis of idiopathic pulmonary fibrosis (IPF). The diagnosis was established according to the American Thoracic Society/European Respiratory Society/Japanese Respiratory Society/Latin American Thoracic Association (ATS/ERS/JRS/ALAT) clinical practice guideline criteria. High-resolution computed tomography (HRCT) scans were required to demonstrate a pattern consistent with usual interstitial pneumonia (UIP) or probable UIP, as defined by the ATS/ERS/JRS/ALAT guidelines.^66^

Blood was collected in tubes and centrifuged at 1500 g for 15 minutes. The top 2/3 of the plasma layer was moved to a new tube and centrifuged again at 1500 g for 15 minutes. A second time, the top 2/3 of the plasma was moved to a new tube to store platelet poor plasma and stored at −80. Samples were thawed and MPO concentration was quantified through ELISA assay based on manufacturer’s instructions.

### Tissue staining for MPO from Human samples

Human formalin fixed paraffin embedded (FFPE) tissue was sectioned into 5 µm slices and stained with anti-MPO (1:100, Goat polyclonal, R&D Systems, AF3667), followed by secondary staining of anti-Goat Alexa Fluor 555 (1:1000) and To-Pro-3 or Hoechst for nuclear localization. Slides were scanned on a Zeiss AxioScan to obtain ∼5-10 mm diameter montage. Image analysis was performed using ImageJ and normalized to the average fluorescent intensity of healthy controls.

### Statistics

Statistical analyses for each experiment are specified in the corresponding figure captions. All analyses were performed using GraphPad Prism v10.6.0. Data distributions were assessed for normality prior to statistical testing. For comparisons between two groups, paired or unpaired two-tailed t-tests were used for normally distributed data, and Mann–Whitney U tests were used when normality assumptions were not met. Comparisons involving three or more groups were analyzed using ordinary one-way ANOVA with Tukey’s multiple-comparisons post hoc test. Experiments involving longitudinal measurements or multiple experimental factors were analyzed using two-way ANOVA or mixed-effects models, as indicated. Survival analyses were performed using Kaplan–Meier methods. Statistical testing was performed only for groups with n ≥ 3. Data are presented as individual biological replicates, as defined in each figure, with summary statistics indicated in the captions. Statistical significance was defined as p < 0.05.

### Data Availability

Values for all data points in graphs are reported in the Supporting Data Values file.

## Results

Given MPO’s role in propagating fibrotic diseases across multiple organs,^67,49–51^ we sought to investigate whether MPO contributes to pulmonary fibrosis development. We studied bleomycin-induced fibrosis in MPOko and WT mice at 14 days post-bleomycin, a time point corresponding to established fibrosis in this model (experimental timeline in Figure 1A). We quantified hydroxyproline content as a measure of total collagen and found MPOko mice were significantly protected from collagen increases after bleomycin administration (Fig 1B). To further test the effects of MPO on fibrosis progression, we treated WT animals with PF1355, a selective MPO inhibitor, daily beginning after the peak in inflammatory cell influx (7 days after bleomycin administration). Like MPOko, mice treated with PF1355 had significantly reduced hydroxyproline levels. Histological analysis reinforced these findings showing both MPOko and MPO inhibition partially protected mice from bleomycin-induced fibrosis across measures of picrosirius staining of collagen content (Fig 1C), lung injury (Supp. Fig 3), and fibrotic area (Supp. Fig 3), with representative images shown (Fig. 1D). However, because MPOko or inhibition only moderately decreased histological findings, MPO inhibition may provide partial, rather than complete, protection against fibrosis.

**Figure 1.**
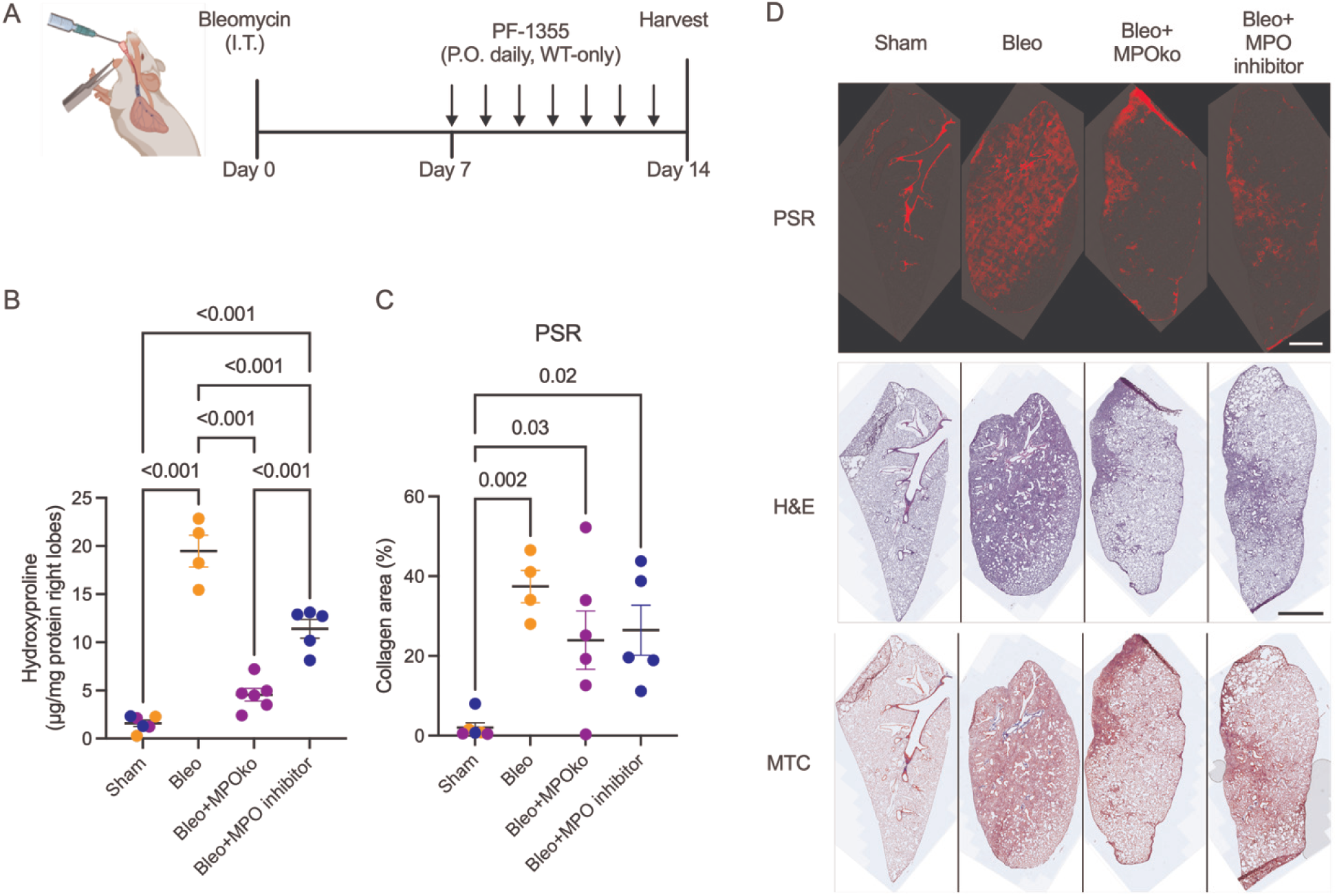
MPOko or MPO inhibition protect against collagen accumulation and fibrotic remodeling after bleomycin injury. A) Experimental timeline showing bleomycin administration, pharmacologic MPO inhibition (PF-1355) beginning at day 7, with tissue harvest at day 14. B) Total collagen content, as measure by hydroxyproline demonstrating significantly reduced collagen accumulation in MPOko mice and in wild-type mice treated with the MPO inhibitor PF-1355 following bleomycin injury. C) Quantification of collagen content from PSR fluorescence showing partial protection from bleomycin-induced fibrosis in MPOko mice and with MPO inhibition D) Representative lung histology from bleomycin-treated wild-type, MPOko, and PF1355-treated mice. N=4-6. Statistical analysis was performed using ordinary one-way ANOVAs with Tukey’s post-hoc test. Each point represents the results from one animal. Bars shown are mean ± SEM.

Because MPO inhibition beginning at day 7 conferred protection against fibrosis, we tested whether MPO influences fibrosis through effects on collagen degradation. CatK has been implicated as a collagen degrading enzyme involved in fibrosis resolution.^19,18,20^ Although CatK expression is increased in bleomycin models and in IPF patient lungs,^19,68^ whether this expression corresponds to increased collagenolytic activity in fibrotic lung tissue in vivo has not been established. Reanalysis of publicly available single-cell RNA-seq data from bleomycin-treated mouse lungs (GSE210341)^60^ confirmed increased *Ctsk* expression after injury, predominantly in adventitial fibroblasts and fibrotic mesenchymal subpopulations (Fig 2A). These data confirm that CatK-expressing cells are present within the fibrotic lung and represent plausible targets for MPO-mediated regulation. We therefore quantified CatK activity in our mouse model. WT mice exhibited reduced CatK activity at day 14 post-bleomycin. In contrast, MPOko mice displayed significantly elevated CatK activity compared to bleomycin-treated WT mice (Fig. 2B; Supp. Fig. 4). PF1355 treatment initiated at day 7 recapitulated this effect. The combination of these results, the beneficial effect of MPO inhibition applied only from day 7 onward, and the elevated CatK activity in MPOko indicate that MPO suppresses CatK activity during fibrosis and may provide a mechanistic explanation for impaired collagen degradation.

**Figure 2.**
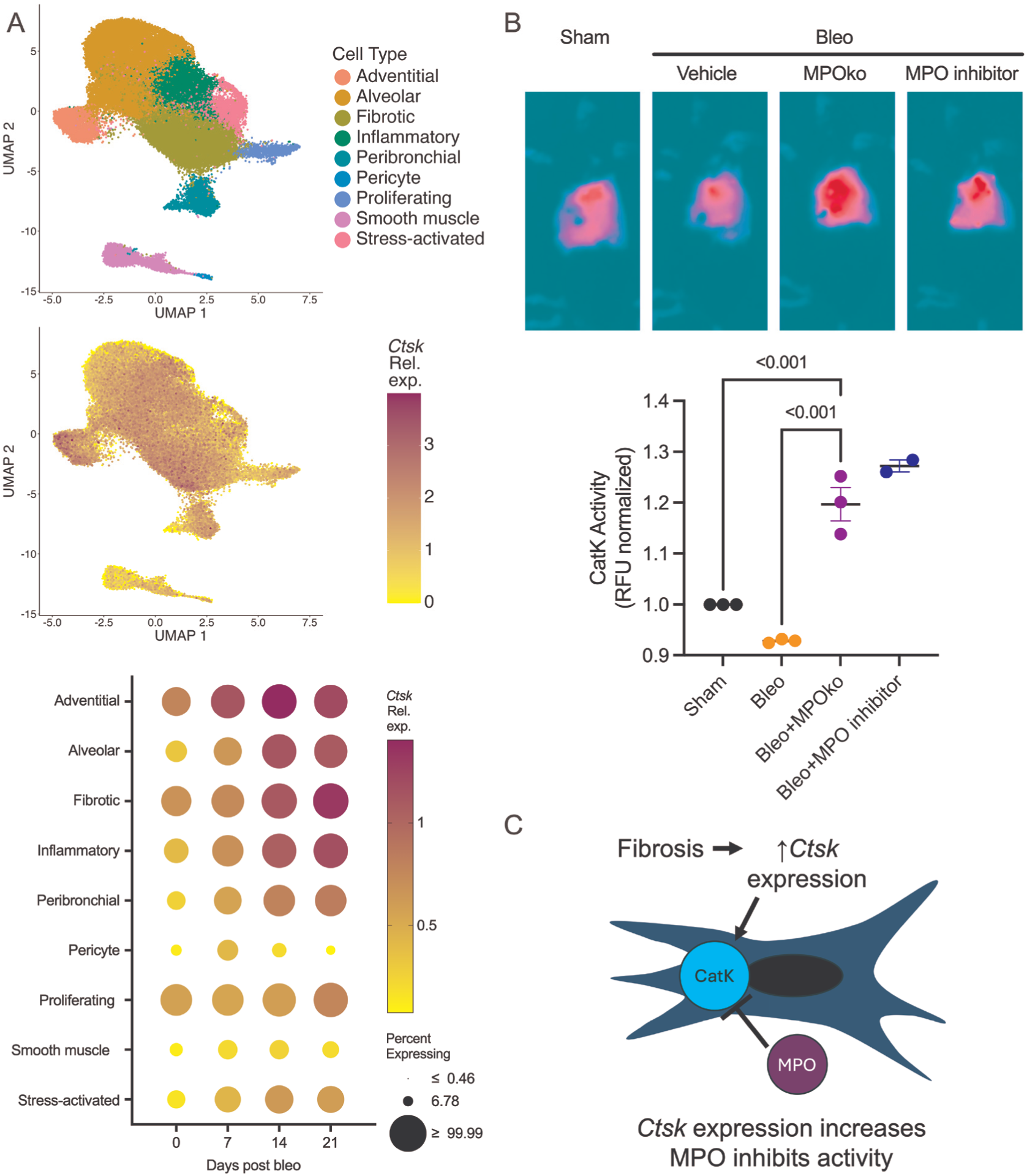
Myeloperoxidase suppresses cathepsin K activity during pulmonary fibrosis in vivo. A) Single-cell RNA-seq analysis of bleomycin-treated mouse lungs showing increased Ctsk expression localized predominantly to adventitial fibroblasts and fibrotic mesenchymal subpopulations (GSE210341 from ^60^). B) CatK activity in lung tissue at day 14 after bleomycin injury, demonstrating reduced activity in wild-type mice and preserved activity in MPOko mice or MPO inhibited mice. C) Conceptual schematic illustrating preserved CatK expression but suppressed CatK activity in the presence of MPO. Together, these data demonstrate uncoupling of CatK expression and activity during fibrosis. Each point represents one independent replicate; n=3; statistical analysis was not performed on the last group because n=2 (all images in Supp Fig 4). Statistical analysis was performed using an ordinary one-way ANOVA with Tukey’s post-hoc test. Bars shown are mean ± SEM.

Because MPO plays well-established roles in acute lung injury, we next investigated whether the protective effects we observed after bleomycin injury reflected altered inflammation or a direct role for MPO in fibrotic tissue. MPOko mice had a higher survival rate (Fig 3A, 94% in MPOko vs 70% in WT) but similar weight changes (Fig. 3B) when compared to wild type controls. MPOko did not affect the abundance of major cell populations during the inflammatory phase of bleomycin-induced fibrosis, including immune, endothelial, epithelial, and tri-lineage–negative populations (Fig. 3C–F; Supp. Fig. 5), indicating that MPO-dependent protection from fibrosis is not attributable to reduced inflammatory cell recruitment.

**Figure 3.**
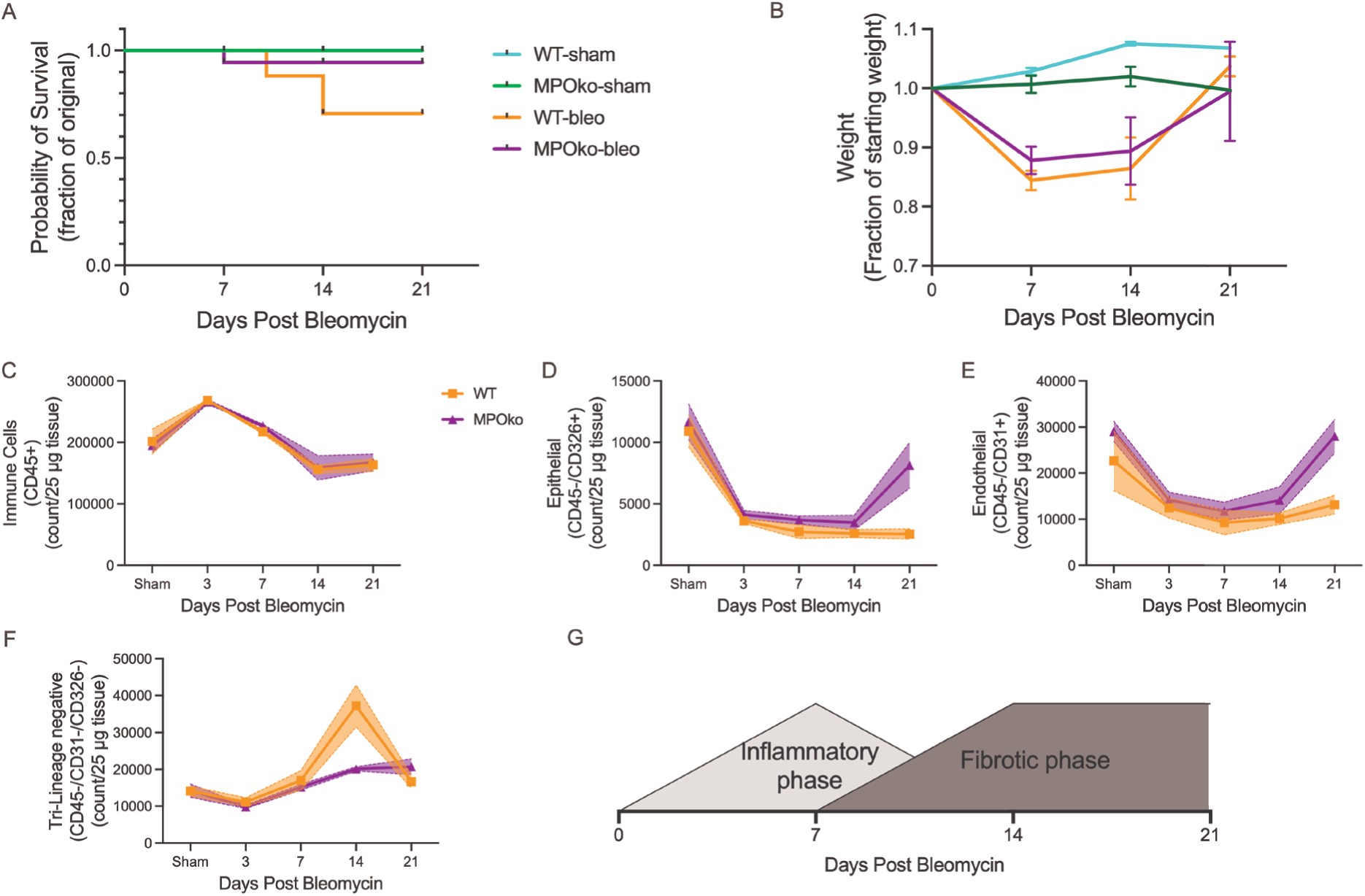
MPO-dependent protection from fibrosis is independent of inflammatory cell recruitment. A) Survival analysis showing improved survival in MPOko mice after bleomycin injury; n=8-20. B) Body weight changes over time demonstrating similar weight loss trajectories between wild-type and MPOko mice; n=3-12. C-F) Flow cytometric quantification of major lung cell populations during the inflammatory phase, including immune (C), endothelial (D), epithelial (E), and tri-lineage–negative cells (F), showing no significant differences between genotypes. G) Schematic delineating the inflammatory and fibrotic phases after bleomycin injury and the timing of MPO inhibition. n=3-4. Statistical analysis was performed using Kaplan Meier survival analysis, or mixed effects models. Each line represents the mean±SEM for each group.

Given that MPOko protected against fibrosis even when MPO inhibition began at day 7 (after peak inflammation), we investigated whether MPO itself persists in lung tissue beyond the inflammatory phase. We first quantified neutrophils (CD45+/Ly6G+/CD11b+) after bleomycin administration in blood, bronchoalveolar lavage fluid (BALF), and tissue. Neutrophils increased transiently after bleomycin administration, peaking in blood at day 3, in BALF at day 7, and in lung tissue at day 14, before returning to baseline across all compartments by day 21 (Fig 4A), consistent with previous reports.^69–72^ We then used ELISAs and immunofluorescence microscopy to quantify MPO levels in tissue after bleomycin administration. Tissue MPO levels were significantly elevated at 7 days and remained significantly elevated at 21 days (Fig 4B) after bleomycin injury. Immunofluorescence staining reinforced these findings, showing MPO levels remained significantly increased 21 days after bleomycin (Fig. 4C). Although neutrophil numbers resolved to baseline levels across all compartments by day 21, MPO protein levels in lung tissue remained significantly elevated through this timepoint. This dissociation between MPO persistence and neutrophil presence suggests that MPO deposited in tissue may exert sustained effects during the fibrotic phase.

**Figure 4.**
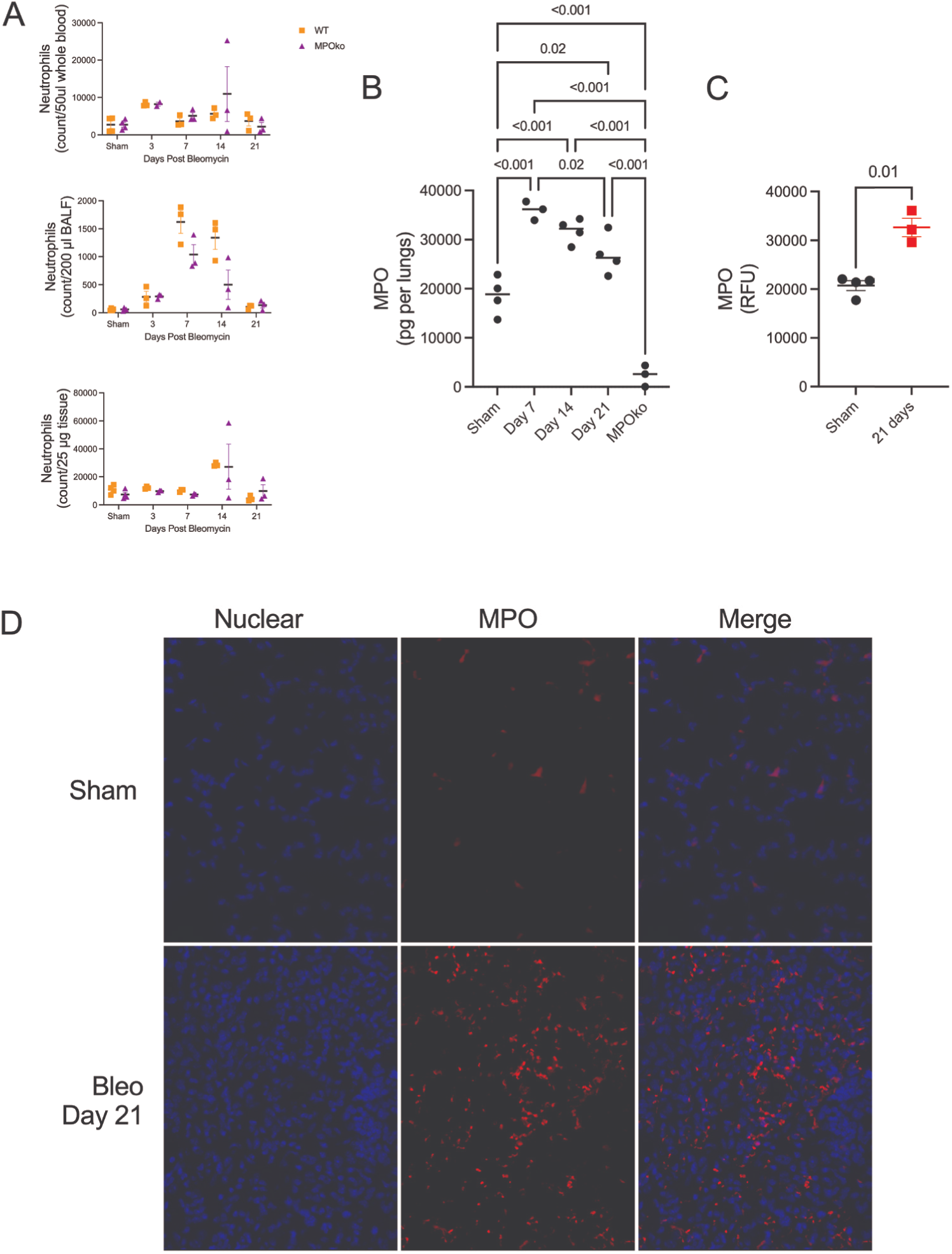
MPO persists in lung tissue beyond neutrophil resolution during fibrosis. A) Neutrophil abundance in blood, bronchoalveolar lavage fluid, and lung tissue over time following bleomycin injury, demonstrating transient infiltration and resolution by day 21, with no difference between genotypes. B) Lung tissue MPO levels measured by ELISA showing sustained MPO presence during the fibrotic phase. C) Immunofluorescent quantification supports persistent MPO elevation at 21 days. N=3-4. Immunofluorescent quantification. D) Statistical analysis was performed using two-way ANOVA mixed effects analysis, Ordinary one-way ANOVA with Tukey’s post-hoc test, or a t-test. Each point represents the results from one animal sample. N=3-4. Bars shown are mean ± SEM.

To determine whether the MPO-dependent suppression of CatK activity observed in vivo reflects a direct effect on CatK activity in lung tissue, we treated precision-cut lung slices (PCLS) with pathophysiologically relevant concentrations of MPO (200 ng/ml from ^73,74^). MPO treatment significantly reduced CatK activity to approximately 50% of control levels, as measured by a fluorescent probe specific for CatK activity (Fig. 5A). Our previous work demonstrated that TGFβ decreases CatK activity,^20^ and we confirmed that finding, with TGF-β treatment decreasing CatK activity to 63% of control levels. Combined treatment of TGF-β and MPO further reduced CatK activity to 38% of control. We next inhibited MPO with ABAH, a selective irreversible MPO inhibitor, which completely attenuated the MPO-induced loss of CatK activity. However, ABAH inhibition of MPO only partially recovered CatK activity back to TGFβ-treated levels in the presence of TGFβ and MPO, reinforcing the in vivo observation that MPO contributes partially, but not exclusively, to impaired collagen degradation. PCLS have also been shown to exhibit spontaneous collagen degrading activity,^75^ so we tested whether pathophysiological levels of MPO would prevent the spontaneous degradation of collagen. To confirm inhibition of CatK activity resulted in decreased collagen degradation, we measured c-terminal cross-linked telopeptide of collagen I (CTX-I), a collagen degradation product specific to CatK activity.^76^ Treatment of precision-cut lung slices with MPO resulted in a ∼20% reduction in CTX-I generation(Fig. 5B) without altering procollagen synthesis (Fig 5C), indicating selective inhibition of collagen degradation rather than synthesis. These findings demonstrate that MPO selectively impairs CatK-mediated collagen degradation rather than collagen synthesis in lung tissue.

**Figure 5.**
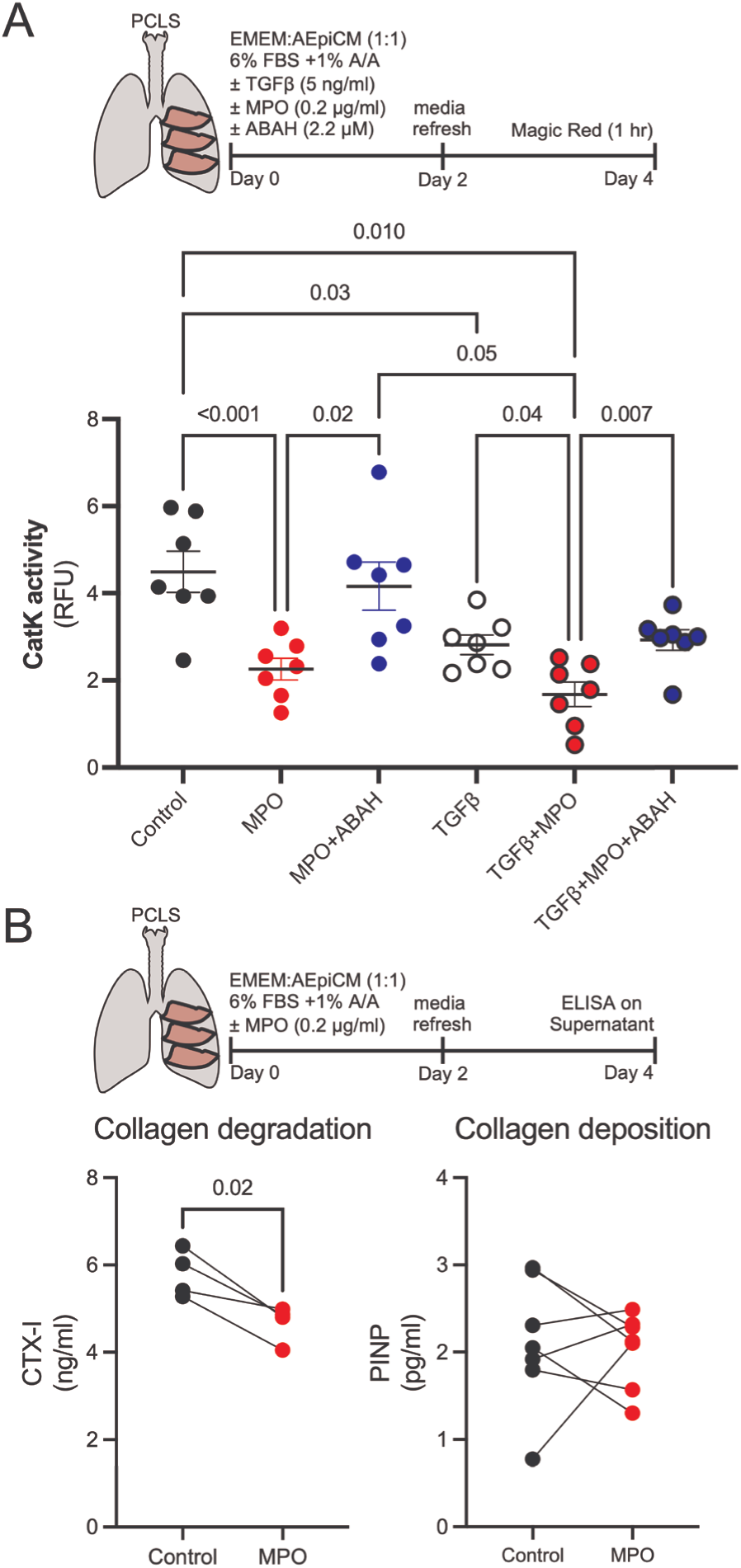
MPO directly inhibits CatK activity and collagen degradation in PCLS. A) CatK activity in PCLS treated with MPO, TGFβ, or both, with or without the MPO inhibitor ABAH. MPO treatment (0.2 µg/ml) decreases CatK activity to 50% of control; combined TGF-β and MPO treatment reduces activity to 38% of control. The MPO inhibitor ABAH restores MPO-induced CatK loss. N=7 Bars shown are mean ± SEM. B) Quantification of collagen degradation (CTX-I) and collagen synthesis (PINP) release from PCLS demonstrating 20% reduced collagen degradation following 4 days of MPO (0.2 µg/ml) treatment. n=4. Statistical analysis was performed using ordinary one-way ANOVA with Tukey’s post-hoc test, or a paired t-test. Each point represents the average results from combining three PCLS from one animal (connected by lines).

Given that MPO persists in tissue and promotes fibrosis independently of inflammation, we sought to identify the mechanism by which MPO inhibits CatK activity. MPO uses H_2_O_2_ to produce HOCl as part of innate immune function, and previous studies have shown that H_2_O_2_ regulates CatK activity directly.^22^ We thus tested both MPO alone and in combination with its substrate, H_2_O_2_, in isolation. MPO directly inhibited recombinant CatK activity in the absence of hydrogen peroxide, reducing activity to 33% of control levels (Fig 6B). Addition of hydrogen peroxide further suppressed CatK activity, and this inhibition occurred at lower hydrogen peroxide concentrations in the presence of MPO (Fig 6C), indicating cooperative regulation. Control experiments confirmed that neither MPO nor hydrogen peroxide affected the fluorescent CatK substrate itself, confirming that the effect was on CatK activity (Supp. Fig 2). These data demonstrate that MPO directly inhibits CatK activity and sensitizes CatK to further inhibition by H_2_O_2_.

**Figure 6.**
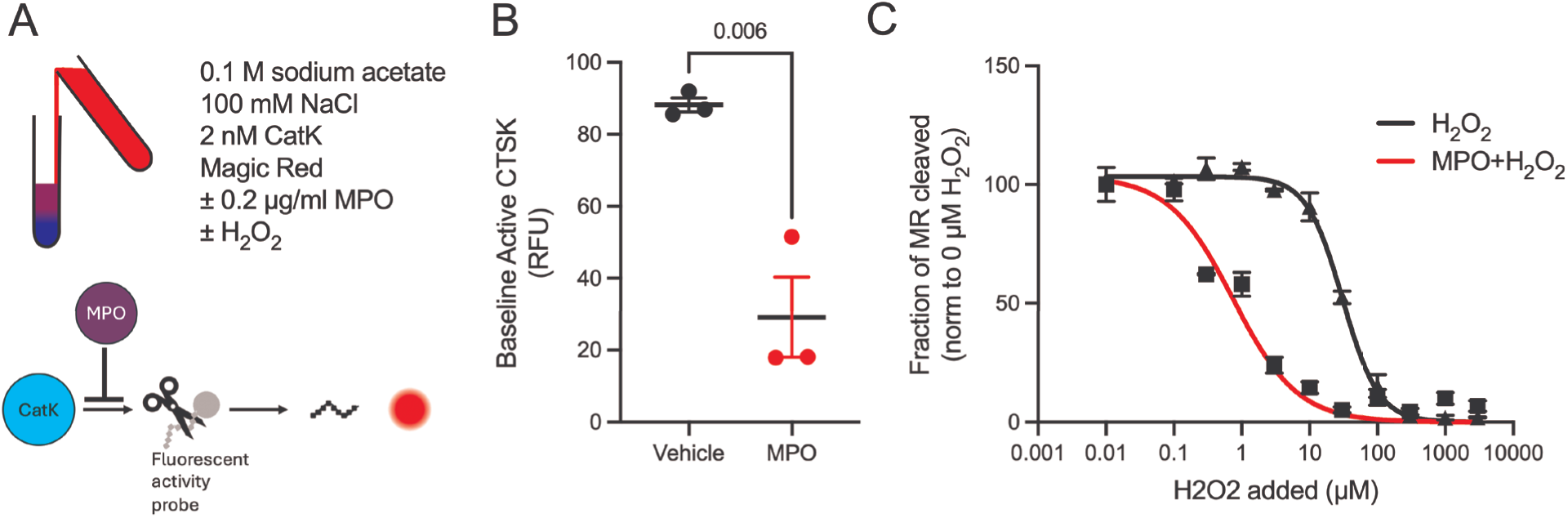
MPO directly inhibits cathepsin K activity. A) Schematic of experimental setup. B) MPO (200 ng/ml) inhibits recombinant CatK activity to 33% of control in the absence of hydrogen peroxide. C) MPO lowers the hydrogen peroxide concentration required to inhibit CatK activity. N=3. Statistical analysis was performed using an unpaired t-test. Each point represents the results from a different MPO source. Bars shown are mean ± SEM.

Having established MPO-dependent suppression of CatK activity in mouse lung tissue, we next examined whether the cellular context required for this mechanism is present in humans. Reanalysis of a publicly available single-cell RNA-seq dataset from healthy human lungs revealed *CTSK* expression enriched in fibroblasts, with highest expression in type 2 alveolar (adventitial) fibroblasts and minimal expression in epithelial, endothelial, or immune populations (Fig. 7A; LMEX0000004396 from ^59^). To determine whether MPO-dependent CatK inhibition occurs in fibroblasts directly, we next measured CatK activity in human lung fibroblasts under profibrotic conditions. Treatment of lung fibroblasts (IMR-90s) with TGFβ alone reduced CatK activity by 20%, consistent with prior reports.^20^ Combined treatment with TGFβ and MPO produced a further significant reduction, reducing CatK activity by 40% (Fig 7B). These findings demonstrate that MPO directly suppresses CatK activity in lung fibroblasts, identifying a specific cellular mechanism through which MPO can impair collagen degradation.

**Figure 7.**
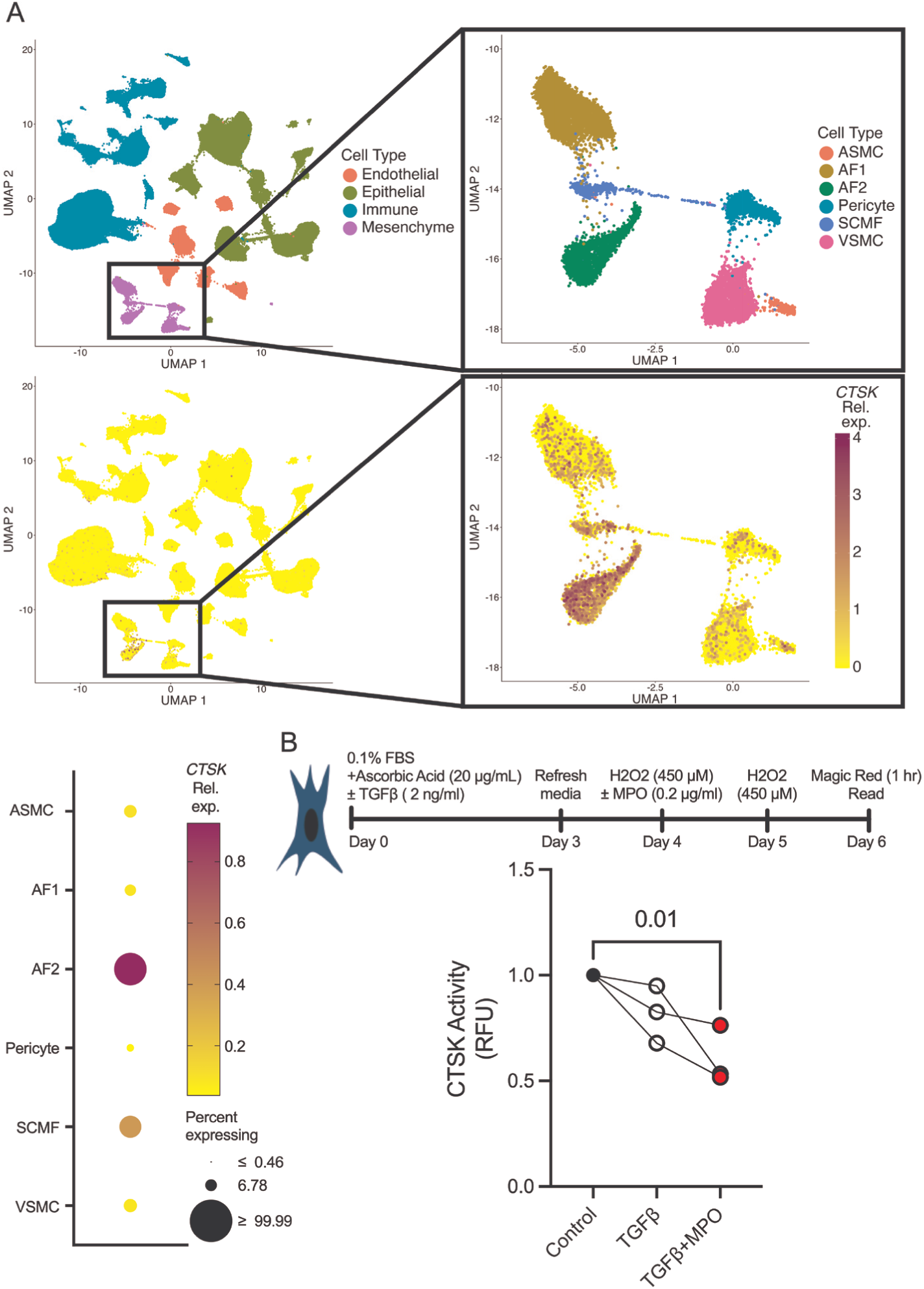
Type 2 alveolar fibroblasts are predominant *CTSK* expressors in healthy lung. A) Reanalysis of previously published Single-cell RNA sequencing of healthy human lungs (LMEX0000004396 from ^59^) identifies AF2 fibroblasts as primary CTSK expressors. B) In IMR-90 human lung fibroblasts, combined TGF-β (2 ng/ml) and MPO (0.2 µg/ml) treatment reduces CatK activity to 60% of control. Statistical analysis was performed using an ordinary one-way ANOVA with Tukey’s post-hoc test. Each point represents one independent replicate N=3.

Because cellular conservation of MPO-mediated CatK inhibition, we next asked whether the cellular and exposure context required for this mechanism is present in human pulmonary fibrosis. Analysis of an independent single-cell RNA-seq dataset comprising healthy control and IPF lungs demonstrated that *CTSK* expression persists in adventitial fibroblasts and is also detected in myofibroblast subpopulations in IPF, with increased average expression and a higher fraction of *CTSK*-expressing cells compared to healthy controls (Fig. 8A; GSE132771 from ^61^). These data indicate that *CTSK*-expressing fibroblast populations are maintained and expanded in fibrotic human lungs, establishing the presence of a potential cellular substrate for MPO-mediated CatK regulation in disease. Immunofluorescent analysis of lung tissue from IPF patients demonstrated increased MPO staining compared to control lungs (Fig. 8B, C). We then assessed whether MPO levels are elevated in patients with pulmonary fibrosis. Consistent with prior reports linking MPO to reduced pulmonary function and mortality in pulmonary fibrosis,^23,31,52^ circulating MPO levels were significantly increased in platelet-poor plasma from IPF patients compared to healthy controls (Fig. 8D), with 68% (13/19) of patients exceeding the maximum value observed in controls. Recently, MPO has been suggested to be a biomarker of disease progression in BALF,^31^ so we sub-grouped the population into MPO-high and MPO-low patients (stratified at 80 ng/mL based on healthy-control maximum) and quantified survival. Elevated MPO levels track with worse clinical outcomes. MPO-high patients had a median survival of 1.5 years, compared to 4.8 years in patients with MPO-low group (Fig. 8E; HR-3.798, 95% CI-1.212 to 11.90, p=0.0285) and both Forced Vital Capacity (FVC; slope −0.1349) and Diffusing Capacity of the Lungs for Carbon Monoxide (DLCO; slope −0.1721) correlated inversely with MPO level (Fig. 8F,G), with the slope DLCO reaching significance. These changes may drive fibrotic progression as seen the decreased life expectancy and decreased pulmonary functions with increasing MPO activity. Collectively, these data indicate that CatK-expressing fibroblast populations coexist with elevated systemic and tissue MPO levels in human pulmonary fibrosis, providing disease-relevant context in which MPO-mediated suppression of CatK activity could contribute to impaired collagen degradation. Thus, a collagenolytic enzyme remains present and poised in the fibrotic lung, but MPO restrains its activity; MPO inhibition may release this brake, restore collagen degradation, and favor matrix resolution.

**Figure 8.**
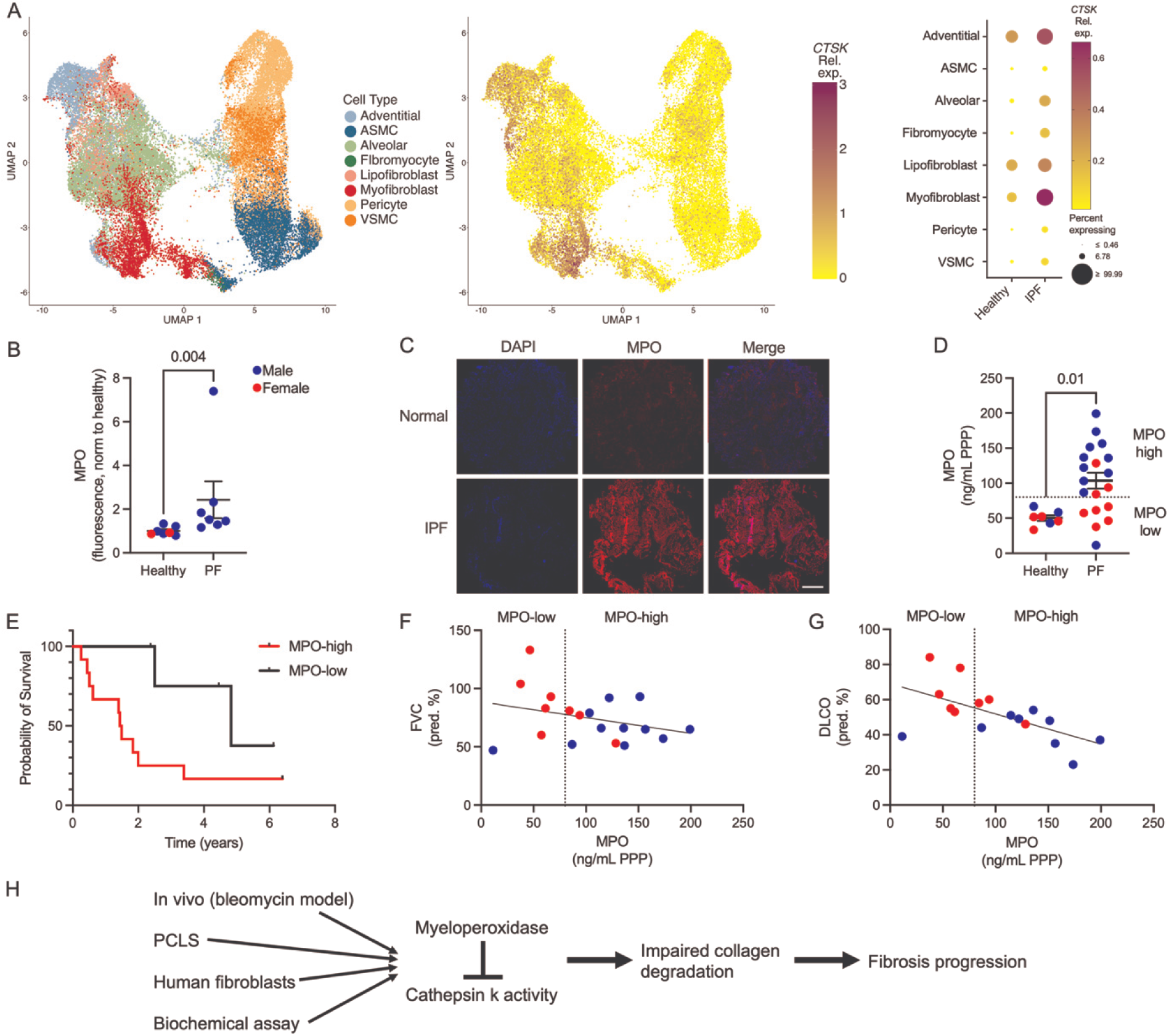
MPO levels are elevated in pulmonary fibrosis patients. A) Reanalysis of single-cell RNA-seq of human lungs (GSE132771 from ^61^) demonstrating *CTSK* expression enriched in fibroblast populations in healthy lungs and increasing in adventitial fibroblasts and myofibroblasts in IPF. B) Lung tissue immunofluorescence shows significantly increased MPO in PF patients. N=7. C) Representative immunofluorescence staining showing increased MPO in fibrotic human lung tissue. D) Circulating MPO is elevated in PF patient platelet poor plasma, with 62% exceeding healthy controls. N=7-19. Patient data was stratified at 80 ng/ml (above healthy controls; dashed line) to establish MPO-high and MPO-low sub-populations. E) MPO-high patients had significantly decreased survival compared with MPO-low patients. MPO-high patients had a median survival of 1.5 years, compared to 4.8 years in patients with MPO-low group (HR-3.798; 95% CI-1.212 to 11.90; p=0.0285). F) Forced Vital Capacity (FVC; slope −0.1349; p=0.20) is inversely correlated with MPO. G) Diffusing Capacity of the Lungs for Carbon Monoxide (DLCO; slope −0.1721, p=0.01) is significantly, inversely correlated with MPO. H) Integrative model summarizing MPO-mediated inhibition of CatK and impaired collagen degradation during fibrosis. Statistical analysis was performed using a Mann-Whitney test (B) because samples failed normality tests, an unpaired t-test (D) for normally distributed values, Wilcoxon test for survival (E), or simple linear regression (F, G). Hazard ratio was calculated using logrank. Each point represents the values from one patient sample. Bars shown are mean ± SEM.

## Discussion

Our results demonstrate that MPO directly diminishes collagen degradation and CatK activity in the lung. MPO treatment of precision-cut lung slices reduced CatK activity and decreased CatK-specific collagen degradation products^16^ while single-cell RNA sequencing identified type 2 alveolar fibroblasts as the predominant CatK expressors in human lungs. Given CatK’s known regulation by oxidants,^16,17,22^ we demonstrated that MPO inhibits CatK directly and independently of H₂O₂, reducing activity to 33% of control. In the bleomycin model, MPOko mice exhibited elevated CatK activity and significant protection from collagen accumulation, effects recapitulated by pharmacological MPO inhibition beginning at day 7 post-injury. This protection occurred independently of effects on inflammatory cell recruitment, and MPO persisted in lung tissue at 21 days despite neutrophil numbers returning to baseline. Among IPF patients, 62% had circulating MPO levels exceeding the maximum healthy control value, with lung tissue staining revealing significantly elevated MPO. Notably, a subset, consisting of over half of PF patients (52%; 10 of 19), exhibited plasma MPO concentrations above 96 ng/mL, a threshold previously associated with increased mortality in other disease contexts.^74^ These findings implicate MPO as a direct inhibitor of collagen degradation that persists beyond the inflammatory phase and may contribute to fibrosis progression in a subset of patients.

These findings fit within an evolving understanding of neutrophil contributions to pulmonary fibrosis. In stable IPF patients, neutrophilia remains mild to moderate, limiting its diagnostic utility,^36^ yet increased fibrotic burden and mortality correlate strongly with neutrophil infiltration.^5,27,77–79^ In acute exacerbations, neutrophil counts rise markedly and predict short-term mortality,^80^ while neutrophil extracellular traps (NETs) accumulate in IPF lungs and correlate with disease severity.^31^ MPO itself predicts faster deterioration of lung function and mortality.^23,31,52^ Our identification of a specific MPO-CatK axis provides mechanistic grounding for these clinical associations.

Previous interventional studies support the therapeutic relevance of targeting neutrophil-derived components over neutrophil presence alone. In the bleomycin model, neutrophil infiltration occurs at early stages and returns to baseline while fibrosis continues.^69–72^ While broad neutrophil depletion in the bleomycin model produces ambiguous results,^37–40^ targeting specific components (neutrophil elastase, PAD4, FPR1) consistently protects against fibrosis.^81–83^ Intratracheal peroxidase administration causes severe lung injury progressing to interstitial fibrosis,^84^ and MPO contributes to fibrosis across multiple organ systems.^49–51^ Tissue-bound MPO persists long after neutrophil resolution,^46–48^ associates with endothelial cells, basement membrane, and glycosaminoglycans, and exhibits increased activity when bound.^85–87^ Multiple groups have identified MPO as a predictor of poor outcomes in pulmonary fibrosis;^23,24,31^ our findings offer CatK inhibition as one mechanism through which persistent accumulation of tissue MPO propagates disease by impairing homeostatic collagen resorption.

MPO represents an attractive therapeutic target because its profibrotic contribution appears modifiable. The specific MPO inhibitors AZD4831 (Mitiperstat) and AZD3421 (Verdiperstat) have demonstrated acceptable safety profiles in clinical trials,^88,89^ although Mitiperstat recently failed primary endpoints in Phase 2/3 trials for metabolic liver disease.^88^ While systemic MPO inhibition carries predictable risks related to impaired antimicrobial oxidant production, the generally mild phenotype of congenital MPO deficiency suggests a therapeutically exploitable safety window, particularly if MPO inhibition is appropriately timed and guided by biomarker status. Identification of a patient subset with elevated MPO levels provides a potential framework for such stratification.

Several limitations warrant acknowledgment. Our human PF cohort remains relatively small (n=19 for plasma, n=7 for tissue), predominated by male samples. Larger studies will be needed to determine whether plasma MPO elevation correlates with disease progression, comorbidities, or treatment response. Additionally, while we present strong functional evidence for MPO inhibiting CatK, MPO affects many other ECM components through crosslinking, protein destabilization, and MMP modulation,^42,41,90–94^ and the relative contribution of CatK inhibition versus these alternative pathways requires further characterization.

In conclusion, we identify MPO as a significant regulator of the balance of collagen synthesis and degradation in lung cells and tissue through its direct inhibition of CatK activity. MPO persists in lung tissue beyond the inflammatory phase, and elevated MPO levels characterize a subset of IPF patients who may benefit from targeted intervention. More broadly, our results highlight the need to study the role of MPO in fibrosis in greater detail and underscore the complexity of targeting collagen degradation globally, raising the likelihood that multiple pathway approaches may be needed to restore normal collagen homeostasis and should prompt detailed evaluation of MPO as a therapeutic target.

## Acknowledgements

The authors acknowledge funding for this project from the American Lung Association Dalsemer ILD Award DAALA2023-1045091 (PAL), Boehringer Ingelheim ILDDA, DoD CDMRP PRMRP HT9425-24-1-0208 (PAL, HN), NIH HL152967, HL166187 (DJT), and HL180318 (YSP). We also acknowledge the C-SiG Clinical Core (DK084567) for use of the Zeiss Axio Observer, and Dr. Kah Whye Peng for the use of the IVIS. The authors acknowledge Claude.Ai for manuscript editing of text.

