## Supplement for "Myeloperoxidase promotes fibrosis by inhibiting cathepsin K to bias the lung toward ECM accumulation"

**Supplemental figure 1. Flow sorting strategy.**

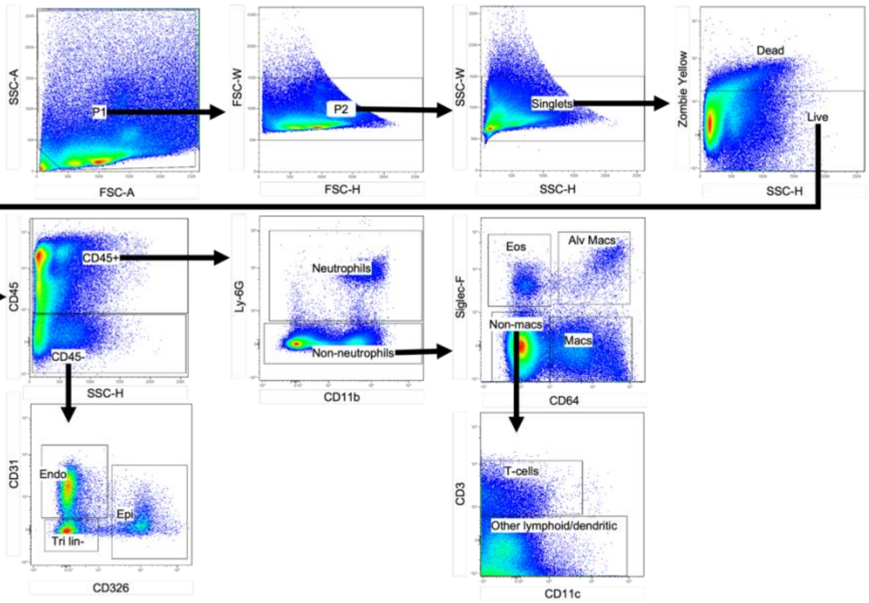

**Supplemental figure 2. Fluorescent probe of cathepsin k activity does not increase in response to H2O2 or MPO.**

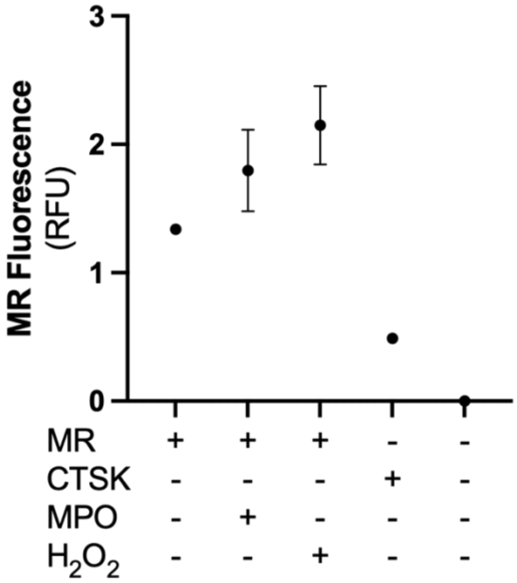

**Supplemental Figure 3. Quantification of fibrotic area analysis from MTC (A) and lung injury from H&E staining (B)**.

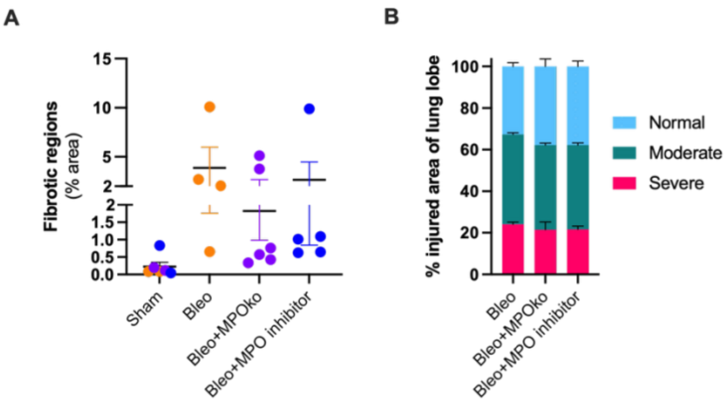

**Supplemental figure 4. IVIS and homogenized lung tissue with cathepsin k fluorescent activity probe demonstrate increased cathepsin k activity in MPOko mice after bleomycin.**

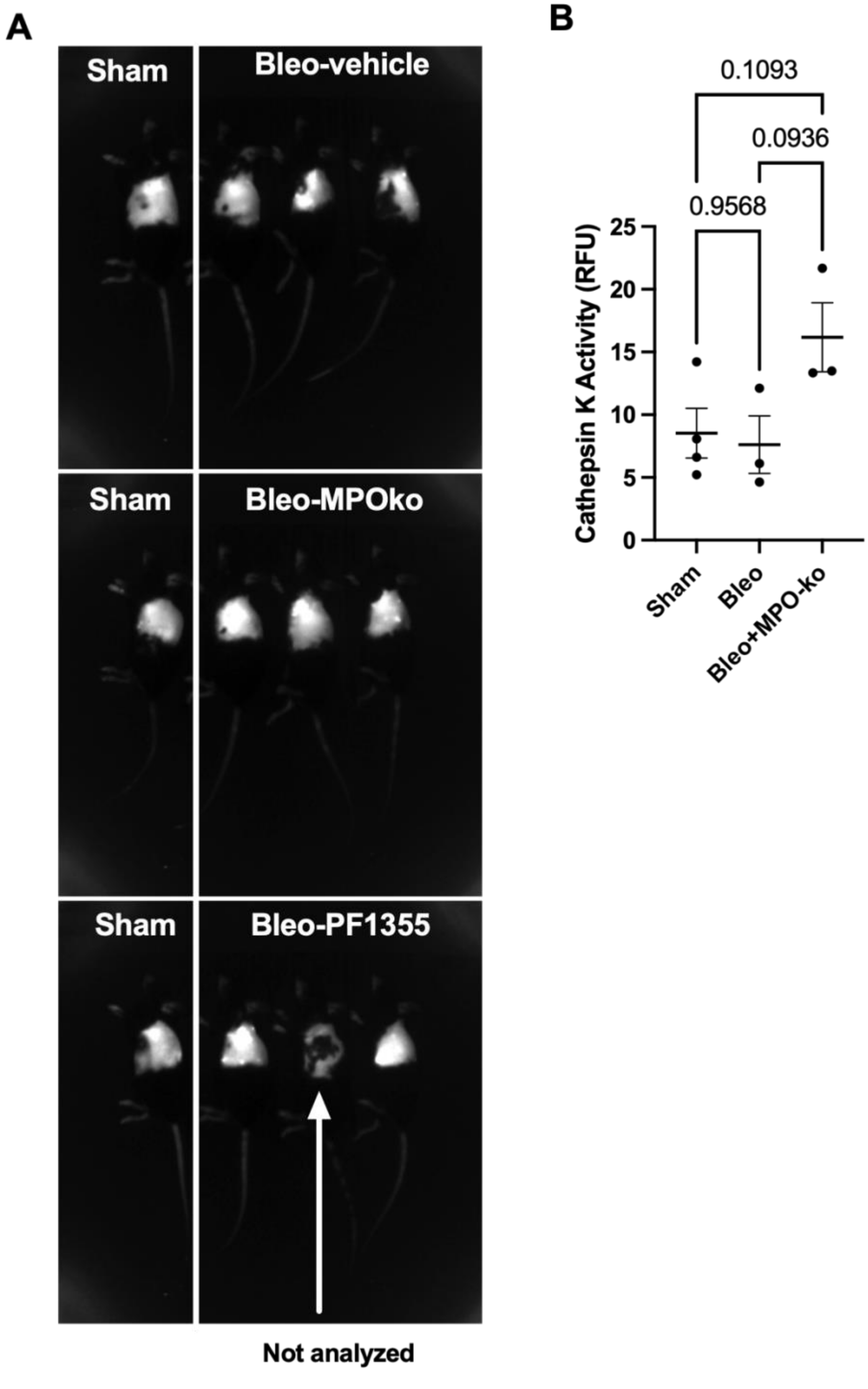

**Supplement figure 5. Frequencies of immune cells are not different between MPOko and WT animals after bleomycin.**

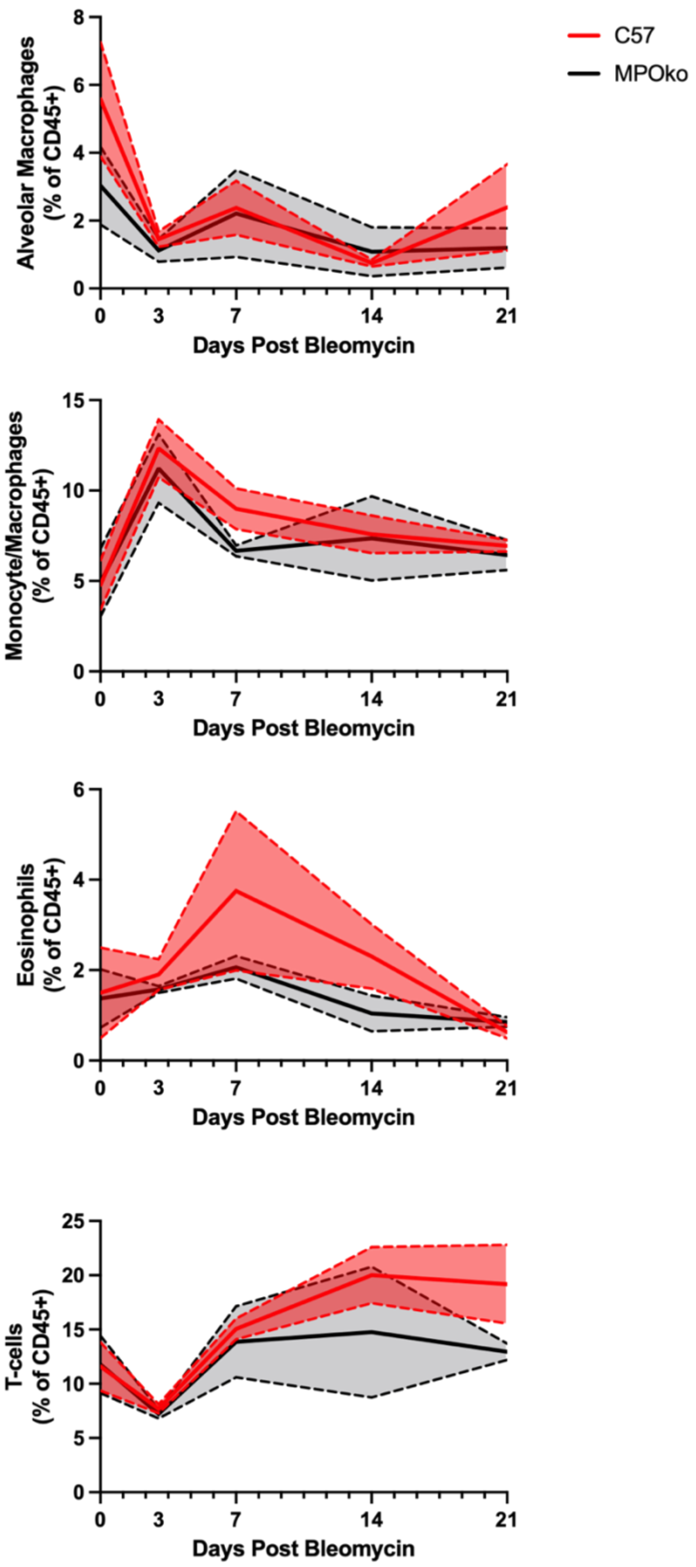

**Supplemental Table 1. Patient demographics for samples used in this study.**

| Patient Demographics | | | | | |
|---|---|---|---|---|---|
| **Characteristics** | | **IPF** | | **Healthy** | |
| Median age (Range) | | 67 (47-86) | | 65 (47-82) | |
| Gender | | | | | |
| | M | 18 | 67% | 9 | 56% |
| | F | 9 | 33% | 7 | 44% |
| CT pattern | | | | | |
| | Probable UIP | 12 | 44% | - | - |
| | UIP | 6 | 22% | - | - |
| | Indeterminate | 1 | 4% | - | - |
| | Unknown | 8 | 30% | - | - |
| Antifibrotic | | | | - | - |
| | Yes | 6 | 22% | - | - |
| | No | 13 | 48% | - | - |
| | Unknown | 8 | 30% | - | - |

**Supplemental table 2. Reagents used in this study.**

| Supplemental Table 2. Equipment and Products Used in These Studies | | | | | |
|---|---|---|---|---|---|
| **Category** | **Product** | **Product Number** | **RRID** | **Vendor** | Model/Version/Lot |
| **Reagents** | Hospira Bleomycin | 0409-0332-20 | — | Vizient Inc. | BL12303B |
| | 20 ga catheter, Angiocath | 26742 | — | BD | |
| | PBS | 14190250 | — | Life Technologies | |
| | PF-1355 MPO inhibitor | HY-100873 | — | MedChemExpress | |
| | Isoflurane | | — | | |
| | Ketamine | | — | | |
| | Xylazine | | — | | |
| | Liberase TM | 5401127001 | — | Sigma-Aldrich | |
| | DNase I | 4536282001 | — | Sigma-Aldrich | |
| | 40 µm filter | SCNY00040 | — | Sigma-Aldrich | |
| | RBC Lysis buffer | 420301 | — | BioLegend | |
| | 30 µm filter | NC9682496 | — | Sysmex America Inc. | |
| | AutoMACS Buffer | 130-091-222 | — | Miltenyi Biotec | |
| | Eagle's Minimum Essential Medium (EMEM) | 30-2003 | — | ATCC | |
| | Alveolar Epithelial Medium Complete Kit (AEpiCM) | 3201 | — | Sciencell | |
| | DMEM | 11965118 | — | Life Technologies | |
| | Anti-Anti | 15-240-062 | — | Fisher Scientific | |
| | FBS | 30-2020 | — | ATCC | |
| | Trypsin-EDTA Solution 0.25% Cell Culture Tested | T4049-100ML | — | Sigma-Aldrich | |
| | Alexa Fluor® 700 anti-mouse CD3 Antibody | 100216 | AB_493697 | BioLegend | |
| | PerCP/Cyanine5.5 anti-mouse CD11c Antibody | 117328 | AB_2129641 | BioLegend | |
| | APC/Cyanine7 anti-mouse/human CD11b Antibody | 101226 | AB_830642 | BioLegend | |
| | PE anti-mouse CD64 (FcγRI) Antibody | 161003 | AB_2904306 | BioLegend | |
| | PE/Cyanine7 anti-mouse Ly-6G Antibody | 127618 | AB_1877261 | BioLegend | |

| | | | | |
|---|---|---|---|---|
| | Brilliant Violet 421™ anti-mouse CD170 (Siglec-F) Antibody | 155509 | AB_2810421 | BioLegend |
| | Zombie Yellow™ Fixable Viability Kit | 423104 | — | BioLegend |
| | BV711 anti-mouse CD45 Antibody | 103147 | AB_2564383 | BioLegend |
| | Brilliant Violet 785™ anti-mouse CD31 Antibody | 102435 | AB_2810334 | BioLegend |
| | Brilliant Violet 650™ anti-mouse Ly-6C Antibody | 128049 | AB_2800630 | BioLegend |
| | FITC anti-mouse CD326 (Ep-CAM) Antibody | 118208 | AB_1134107 | BioLegend |
| | Purified Anti-Mouse CD16 / CD32 (Fc Shield) (2.4G2) | 70-0161-U500 | AB_2621487 | Cytek |
| | True-Stain Monocyte Blocker | 426102 | — | BioLegend |
| | Masson's Trichrome Stain | 25088-1 | — | Polysciences Inc. |
| | Hematoxylin Solution, Gill No. 3 | GHS332-1L | — | Sigma-Aldrich |
| | Eosin Y | E4009 | — | Sigma-Aldrich |
| | Picro-Sirius Red Solution | ab246832 | — | Abcam Inc. |
| | To-Pro-3 | T3605 | — | ThermoFisher |
| | Hoechst 33342, trihydrochloride trihydrate | H1399 | — | ThermoFisher |
| | Alexa 555 anti-goat secondary antibody | 62248 | — | ThermoFisher |
| | Hydroxyproline Colorimetric Assay Kit | K555-100 | — | BioVision |
| | PINP ELISA kit: Mouse PINP ELISA Kit | MBS2500076 | — | MyBioSource |
| | Human Myeloperoxidase ELISA Kit - Quantikine | DMYE00B | — | R&D Systems |

| | | | | |
|---|---|---|---|---|
| | CTXI ELISA kit: Mouse CTXI (Cross Linked C-Telopeptide of Type I Collagen) ELISA Kit | MBS9141384 | — | MyBioSource |
| | MPO ELISA DuoSet anti-Mouse | DY3667 | — | R&D Systems |
| | UltraPure™ Low Melting Point Agarose | 16520100 | — | ThermoFisher |
| | Gelatin from porcine skin, powder, gel strength ~300 g Bloom, Type A, BioReagent, for electrophoresis, suitable for cell culture | G1890-100G | — | Sigma-Aldrich |
| | CaCl2 | C4901-500G | — | Sigma-Aldrich |
| | MgSO4 | M7506-500G | — | Sigma-Aldrich |
| | Magic Red | 940 | — | ImmunoChemistry |
| | recombinant active MPO | 3174-MP-250 | — | R&D |
| | MPO isolated from human leukocytes | 475911 | — | Sigma-Aldrich |
| | MPO isolated from human leukocytes | M6908 | — | Sigma-Aldrich |
| | Recombinant Active Human Cathepsin K | ab157067 | — | Abcam |
| | In vivo cathepsin K activity reporter | NEV11000 | — | Perkin Elmer |
| | anti-MPO antibody | AF3667 | AB_2250866 | R&D Systems |
| | 4-Aminobenzoic Acid hydrazide (ABAH) | 14845 | — | Cayman Chemical |
| | Human TGF-beta 1 Recombinant Protein, PeproTech® | 100-21C-10UG | — | ThermoFisher |
| | Hydrogen Peroxide | 16911-250ML-F | — | Sigma-Aldrich |
| | EDTA, 0.5M Solution 100ml | C001N18 - 4055-100ML | — | Thomas Scientific |
| | T-PER™ Tissue Protein Extraction Reagent | 78510 | — | ThermoFisher |

| | | | | | |
|---|---|---|---|---|---|
| | HBSS (10x), no calcium, no magnesium, no phenol red | 14185052 | — | ThermoFisher | |
| | Sodium Acetate, 3M, pH 5.2, Molecular Biology Grade - CAS 127-09-3 - Calbiochem | 567422 | — | Sigma-Aldrich | |
| | L-Ascorbic Acid (White Crystalline Powder), Fisher BioReagents | BP351-500 | — | Fisher Scientific | |
| **Organisms** | IMR-90 (human lung fibroblast) | CCL-186 | CVCL_0347 | ATCC | |
| | MPO-ko mice (B6.129X1-Mpotm1Lus/J) | 004265 | IMSR_JAX:004265 | Jackson Labs | |
| | C57BL/6J mice | 000664 | IMSR_JAX:000664 | Jackson Labs | |
| **Equipment** | LSR Flow Cytometer | Fortessa X-20 | — | BD Biosciences | |
| | Fluorescent Plate Reader | FlexStation 3 | — | Molecular Devices | |
| | Brightfield Slide Scanner | EasyScan Pro | — | Motic | |
| | Axio Observer Widefield Fluorescent Microscope | Axio Observer | — | Ziess | |
| | Tissue Homogenizer | BeadMill 24 | — | ThermoFisher | |
| | Axioscan slide scanner | AxioScan | — | Zeiss | |
| | IVIS Spectrum In Vivo Imaging Systems | IVIS Spectrum | — | Perkin Elmer | |
| | Microscope (Confocal) | CKX53 | — | Olympus | |
| **Software** | EasyScan | | SCR_024854 | Motic | |
| | Statistical Analysis Software | | SCR_002798 | GraphPad Prism | v10.6 |
| | Image Analysis Software | | SCR_002285 | ImageJ (Fiji) | v1.53t |
| | Flow Cytometry Analysis Software | | SCR_008520 | FlowJo | v10.8 |
| | Fluorescent Plate Reader Software | | SCR_014240 | Molecular Devices SoftMax Pro | v7 |
